# Stretching drives Membrane Homogenization in Phase-Separated Supported Lipid Bilayers

**DOI:** 10.64898/2026.02.26.708304

**Authors:** Aviya Perlman Illouz, Ruth Meyer, Sarah Köster, Gonen Golani, Raya Sorkin

**Affiliations:** School of Chemistry, Faculty of Exact Sciences, Tel Aviv University, Tel Aviv, Israel; Center for Physics and Chemistry of Living Systems, Tel Aviv University, Tel Aviv, Israel; Institute for X-Ray Physics, University of Göttingen, Germany; Department of Physics, University of Haifa, 3498838 Haifa, Israel

## Abstract

Cell plasma membranes exhibit heterogeneous lateral organization whose dynamic compartmentalization is critical for processes such as viral infection and fertilization. While membrane tension is known to influence crucial cell remodeling processes, its role in regulating membrane heterogeneous organization remains unclear. To reveal the effect of tension on lateral membrane organization, we used supported lipid bilayers on flexible substrates. These were prepared by rupturing ternary-composition giant unilamellar vesicles exhibiting liquid order-disorder phase coexistence. The phase coexistence is observed using a fluorescent probe that preferentially partitions to the disordered phase. Using a motorized equibiaxial stretching device, we observed progressive homogenization of domain morphology with increasing strain up to a critical strain, beyond which the membrane remains uniform. We define an order parameter based on the relative concentration of the dye in the two phases, which is a proxy for the membrane lateral organization. Order parameter analysis revealed power-law scaling below the critical strain with an exponent β = 1.0 ± 0.3, consistent with an elastic theoretical model predicting β = 1. The progressive broadening of the interfacial region width near the critical strain, and continuous transition to a homogeneous phase, is consistent with a second-order phase transition. These findings indicate that membrane tension may serve as a physical regulator of lateral lipid organization, with implications for how cells use mechanical forces to regulate their structure and function.

**Significance Statement:** Cell membranes are fundamental structures serving as the interface between a cell’s interior and its external environment. Membrane tension regulates critical cellular processes, yet how it affects the essential heterogeneous organization of membrane domains remains controversial. We combine artificial supported membranes on flexible substrates with a motorized equibiaxial stretching device, to directly visualize domain morphology under controlled tension. We observed that stretching drives the membrane to become progressively more uniformly distributed until it appears homogeneous. This transformation is consistent with a second-order phase transition. These findings demonstrate that mechanical tension acts as a direct physical regulator of cell membrane structure and lateral lipid organization.

## Introduction

The dynamic arrangement of phospholipid bilayers with integrated membrane proteins determines essential cellular plasma membrane properties like thickness, fluidity (1), viscosity (2), permeability, and biological functionality (3). Distinct domains arise from lipid–lipid interactions, proteins recruiting specific lipids, or lipid–protein interactions (4, 5). This spatial compartmentalization is crucial for membrane functions, including directing fusion events (6–8), cellular signalling, and trafficking (9). In membrane model systems composed of saturated lipids, unsaturated lipids, and cholesterol, lipids undergo spontaneous lateral organization into distinct liquid phases. This results in cholesterol-enriched liquid-ordered (Lo) domains that coexist with liquid-disordered (Ld) domains.

Membrane tension regulates critical cellular processes, including vesicular trafficking, cell migration, and membrane remodelling (10). It arises from multiple sources - osmotic pressure differences, adhesive interactions with extracellular substrates and neighboring cells, and mechanical forces transmitted through the cytoskeletal network. Tension varies considerably between cell types and throughout the cellular life cycle (11).

Experimental evidence indicates that tension regulates membrane lateral organization (12–16). However, studies examining the interplay between tension and domain formation have yielded contradictory findings, with results depending on the method of tension application. In unilamellar vesicles, tension increase induced by osmotic pressure promotes domain formation by raising the miscibility temperature (12–14). In contrast, tension increase through micropipette aspiration suppresses domain formation by lowering this temperature (15, 16), consistent with theoretical predictions for lipid bilayers (17, 18).

The phase transition between Ld-Lo coexistence and a homogeneous phase can be characterized by an order parameter that follows power-law behavior near the critical point and vanishes at the phase transition. First-order transitions exhibit a discontinuous first derivative of the free energy at the transition point, whereas second-order transitions are continuous (19). The phase behavior of lipid mixtures as a function of temperature has been systematically characterized (20), and the temperature-driven transition from Ld-Lo coexistence to a homogeneous liquid phase was classified as second-order (21). While such thermal phase transitions in lipid membranes are well-established, how tension drives analogous phase transitions remains controversial (12–16).

Here, we use supported lipid bilayers (SLBs) on flexible substrates combined with a motorized equibiaxial stretching device to directly visualize domain morphology under controlled, uniform membrane stretching. The SLBs were prepared by rupturing ternary-composition Giant Unilamellar Vesicles (GUVs) exhibiting Ld-Lo phase coexistence. The phase coexistence is observed using a fluorescent probe, Rhodamine-Phosphatidylethanolamine (Rh-PE), which preferentially partitions to the disordered phase (22). We find that Rh-PE, initially enriched in the Ld phase, becomes progressively more uniformly distributed as strain increases up to a critical threshold, beyond which the membrane appears homogeneous. Concomitantly, the interfacial regions between coexisting phases broaden with increasing strain. The continuous loss of phase coexistence together with diverging interfacial width fluctuations is consistent with a second-order phase transition.

## Results

### Preparation of phase-separated SLBs on flexible substrates compatible with equibiaxial stretching devices

We prepared GUVs composed of DSPC/DOPC/cholesterol (27:50:23) exhibiting clear phase separation at room temperature (Fig. 1A). The membranes were labelled using the Rh-PE, providing clear visualization of domain organization and enabling quantitative analysis of domain properties. SLBs were subsequently created by rupturing the GUVs (Fig. 1B) on flexible PDMS substrates, following an optimized plasma treatment to ensure proper membrane adhesion. During rupture, individual vesicles formed distinct membrane patches on the PDMS surface, rather than forming a continuous bilayer. The resulting SLBs preserved the domain structures as shown in Fig. 1C. We fabricated PDMS samples of a designated geometry, compatible with a custom-designed motorized equibiaxial stretching device (SI Fig. S1).

**Fig. 1:**
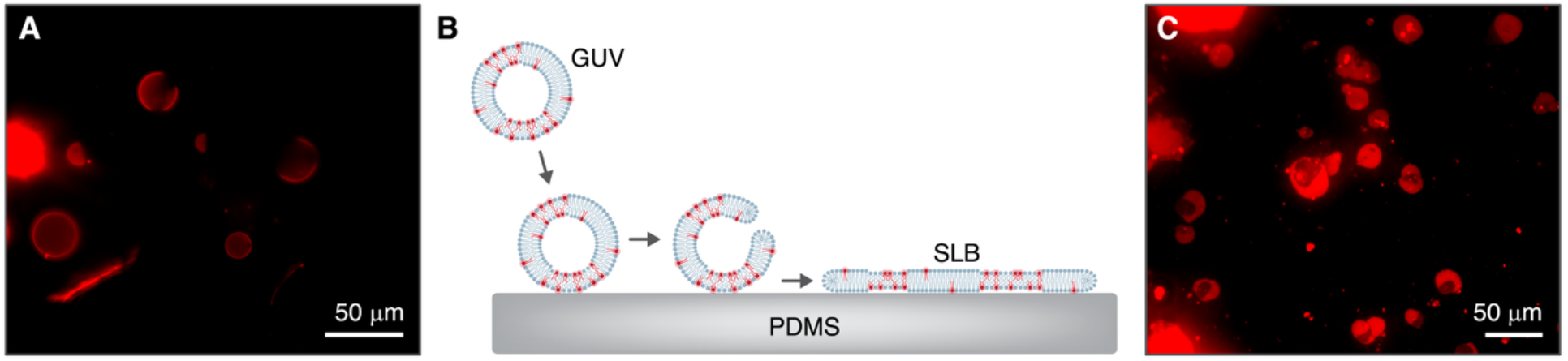
Formation of phase-separated SLBs on stretchable substrates for an equibiaxial stretching device. **(A)** Fluorescence microscopy image of GUVs exhibiting lipid phase separation. The Ld phase appears bright due to preferential partitioning of Rh-PE fluorescent label into this phase, while Lo domains appear dark. GUVs composed of DSPC/DOPC/cholesterol (27:50:23) incorporating 0.1% Rh-PE. **(B)** Schematic illustration of SLB formation through GUV rupture on PDMS substrate (Created in BioRender. PI, A. (2026) https://BioRender.com/2svkach). **(C)** Fluorescence microscopy image of phase-separated SLBs preserve lipid

### Atomic force microscopy (AFM) validates phase separation and reveals a nanoscale height difference between Ld and Lo domains

High-resolution AFM imaging provided direct topographical characterization of phase-separated SLBs, revealing nanoscale structural information for both fluorescently-highlighted bright regions (Ld) and unlabelled dark regions (Lo). Fig. 2A shows a representative fluorescence microscopy image and corresponding AFM topographical scan. A magnified view of the AFM scan (Fig. 2B) revealed distinct height differences between regions corresponding to Ld and Lo domains. Height profiles indicated a total membrane thickness of approximately 4 nm, consistent with phospholipid bilayer dimensions (1) (Fig. 2C). Within the membrane, the Lo domain height was approximately 1 nm higher than that of the surrounding Ld regions, as shown by the profile along the green line (Fig. 2D). This height difference is consistent with the extended, tightly-packed conformation of saturated lipids and cholesterol in ordered phases (5) and agrees with previous studies (23).

**Fig. 2:**
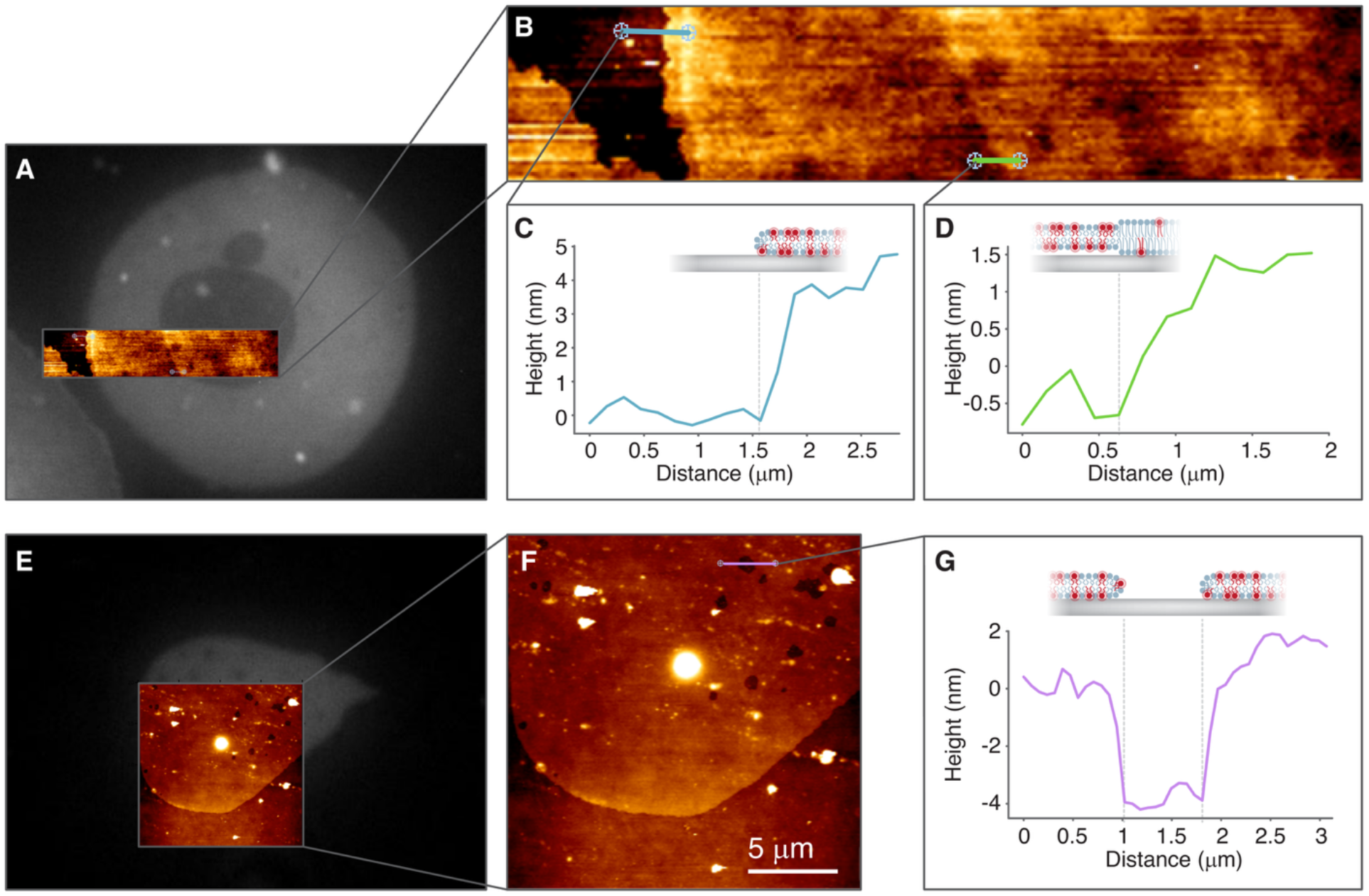
Nano-scale characterization and discrimination of SLBs regions using AFM. **(A)** Correlative fluorescence and AFM imaging of an SLB showing phase separation. **(B)** Magnified AFM scan showing height difference between Ld and Lo domains. **(C)** Height profile along the blue line in panel B, showing ∼4 nm of total bilayer thickness. **(D)** Height profile along the green line in panel B, showing ∼1 nm of height difference between coexisting phase domains. **(E)** Correlative fluorescence and AFM imaging showing membrane defects (holes). **(F)** Magnified AFM scan of membrane defects. **(G)** Height profile across defect showing depth of ∼4 nm, corresponding to the full bilayer thickness, enabling discrimination between holes and Lo domains.

Additional AFM scans (Fig. 2E) examined membrane features that appeared darker than Lo domains in the corresponding fluorescence microscopy. Magnified views and height profiles revealed depths of approximately 4–5 nm, corresponding to the full bilayer thickness (Fig. 2F-G). These measurements confirmed that these darker regions were membrane defects (holes) rather than Lo domains, enabling discrimination between the two dark-appearing features.

The correlation between fluorescence microscopy and AFM scans validated the identification of three distinct membrane regions: high fluorescence Ld domains (bright), low fluorescence Lo domains (dark, elevated by ∼1 nm), and membrane defects (darker, ∼4–5 nm deep). This confirmation of phase-separated domains enabled subsequent quantitative analysis based on fluorescence imaging.

### Increasing area strain induces membrane homogenization

To examine tension-mediated membrane reorganization, SLBs were subjected to controlled mechanical strain using a motorized equibiaxial stretching device. Because the membrane is attached to the substrate, stretching the substrate also stretches the bilayer, thereby increasing membrane tension. The imposed area strain, denoted by *ϵ*, was quantified as described in Methods. The stretching protocol achieved a progressive membrane area strain increase of up to ∼25% (SI Fig. S2).

Representative microscopy images at sequential strain levels are shown in Fig. 3. The complete experimental series, including additional strain levels, is shown in Fig. S4. At zero imposed strain, the membrane displayed a heterogeneous structure characterized by distinct micro-domains with dark Lo phase regions exhibiting minimal Rh-PE fluorescence and bright Ld phase regions (Fig. 3-I). As substrate strain increased (Fig. 3-II), domain boundaries became less sharp, and the fluorescence intensity difference between phases decreased, reflecting partial redistribution of the Rh-PE probe from the Ld phase toward the Lo phase and indicating progressive lipid mixing of the two phases. At a high area strain (Fig. 3-III), domains were no longer visually distinguishable, with fluorescence appearing nearly uniform across the membrane, consistent with redistribution of the Rh-PE probe and transition to a homogeneous membrane state.

**Fig. 3:**
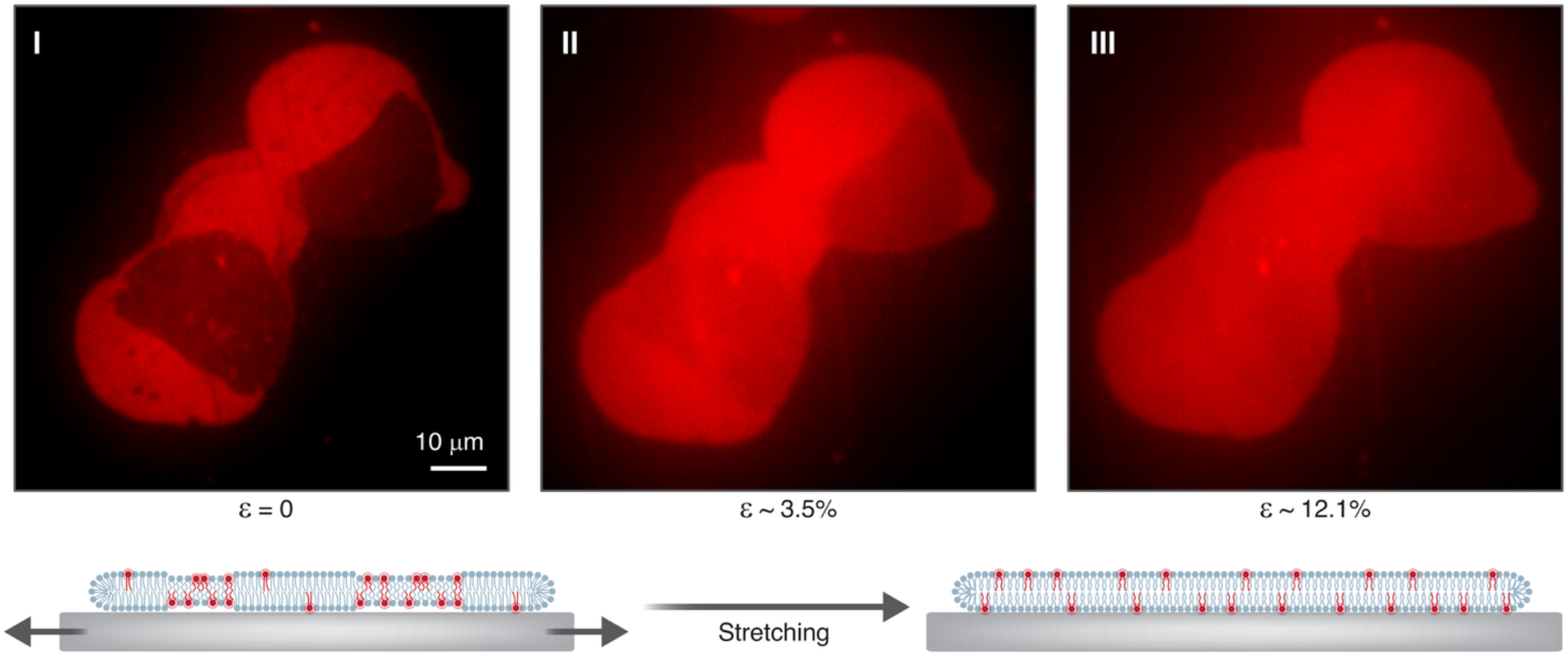
Tension-induced reorganization of membrane domains. Sequential fluorescence images of phase-separated SLBs at increasing stretching levels, demonstrating progressive membrane homogenization as Rh-PE fluorescent probe redistributes from the Ld toward the Lo phase with increasing membrane tension. Images were normalized to account for photobleaching effects during sequential imaging.

Membrane holes defects appeared at higher strain levels (SI Fig. S3), providing direct evidence of mechanical coupling between the bilayer and substrate. The exact point of holes formation in our system is difficult to determine due their small size.

### Membrane homogenization under strain exhibits continuous phase transition behavior with a defined critical point

To quantify membrane homogenization under imposed tension, the Rh-PE fluorescence intensity was evaluated within selected regions of the Lo (dark) and Ld (bright) phases, denoted by *I*_*Lo*_ and *I*_*Ld*_, respectively, and their intensity ratio, *I*_*Lo*_/*I*_*Ld*_, was calculated, as described in Methods. A region-based approach was chosen over area-fraction analysis from whole-membrane intensity distributions because large-area measurements were affected by optical artifacts from particulates in the buffer.

For the representative case shown in Fig. 4A, the intensity ratio began at a low value in the unstretched state, indicating clear phase separation. As area strain increased, the ratio increased progressively, reaching a critical threshold (determined as described in Methods). Beyond this point, the ratio plateaued at a value approaching close to unity, indicating transition to a homogeneous state, though a slight deviation implies that some residual partitioning remains. The data demonstrate characteristic power-law scaling with distinct regimes before and after a well-defined critical strain.

**Fig. 4:**
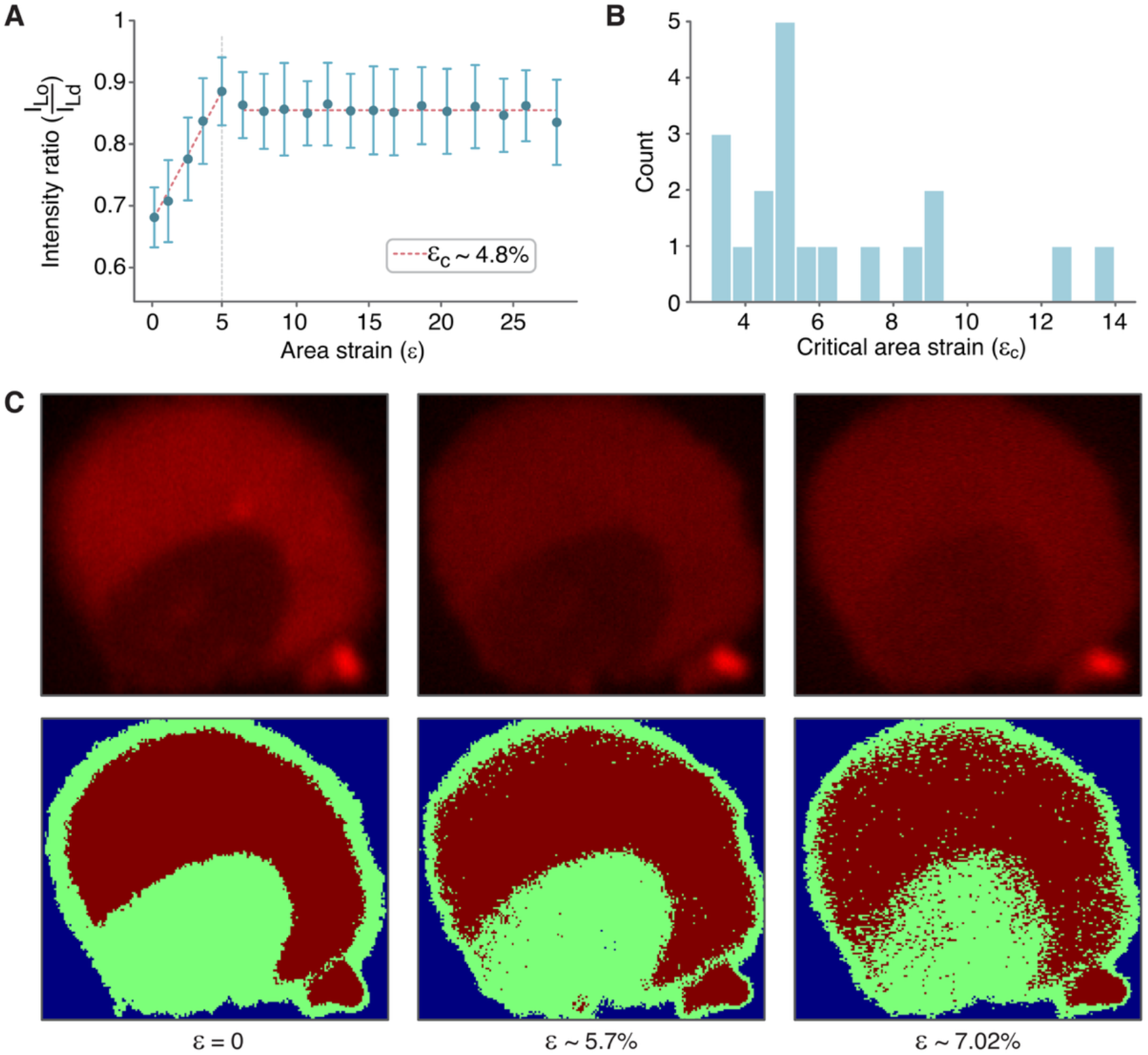
Domain reorganization in SLBs exhibits a phase transition behavior under applied strain. **(A)** Quantification of membrane homogenization vs. area strain, using fluorescence intensity ratio, performed using a region-based approach in which mean intensities were measured within selected regions of interest defined within the Lo and Ld phases. Error bars represent the standard deviation of the mean values across the selected regions. The dashed lines indicate distinct scaling behaviors before and after a well-defined critical strain point, which are consistent with continuous phase transition behavior. **(B)** Distribution of critical strain values across all independent membranes (n=19). **(C)** Segmentation of sequential fluorescence images of a representative membrane using Multi-Otsu thresholding, to classify pixels into Lo and Ld phases at increasing stretching levels. Domain boundaries became progressively less distinct and the interfacial boundary region broadened with increasing membrane strain.

Critical behavior was observed in most of the vesicles examined, indicating that experimental conditions were set close to the critical strain. Analysis of critical strain values across all vesicles yielded a distribution with a mean of *∈*_*c*_ = 6 ± 3% (Fig. 4B).

To further characterize the reorganization associated with the transition, sequential fluorescence images were segmented using Multi-Otsu thresholding to classify pixels into Lo and Ld phases for area fraction estimation at different stretching levels. Quantitative analysis of absolute areas was constrained by incomplete membrane imaging and residual buffer fluorescence, and the total Lo/Ld area ratio could not be tracked with sufficient accuracy across stretch levels (see SI Fig. S5C). However, the segmented images clearly show that domain boundaries became progressively less distinct and the interfacial boundary region broadened with increasing membrane tension. A representative example is shown in Fig. 4C. The progressive increase in Rh-PE intensity ratio and broadening of domain boundaries are consistent with a second order phase transition (19, 24).

### Scaling Law Behavior Near Critical Strain

To analyze the observed transition behavior near the critical strain, an appropriate *order parameter* was defined based on the Rh-PE fluorescence intensity ratio between the Lo and Ld domains. Since Rh-PE preferentially partitions into the Ld phase due to its bulky headgroup (22), this ratio serves as a natural proxy for the degree of membrane heterogeneity. Assuming free exchange of Rh-PE molecules between the Ld and Lo domains, their chemical potentials are equal at equilibrium, and their concentration ratio satisfies:

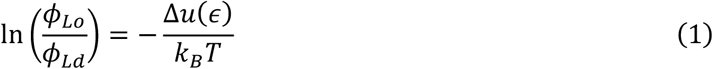

Where Δ*u*(*ϵ*) is the energy difference per Rh-PE molecule between the Ld and Lo phases, *ϕ*_*Lo*_ and *ϕ*_*Ld*_ are the Rh-PE area fractions in the Lo and Ld phases, respectively. We assumed the Rh-PE area fraction to be small in both phases.

Expanding the energy difference near the critical strain, *ε*_*C*_, to leading order, we obtained (SI Section 4) a general form to the energy difference:

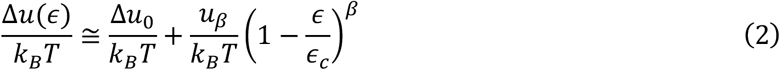

Where *β* is the power of the leading term in the expansion, *u*_*β*_ is its corresponding coefficient, and Δ*u*_0_ is the strain-independent energy difference, arising from the difference in lipid composition between the two domains and from packing constraints in the different phases that govern Rh-PE interaction with its local environment.

Subtracting the concentration ratio at the critical strain yielded the following order parameter:

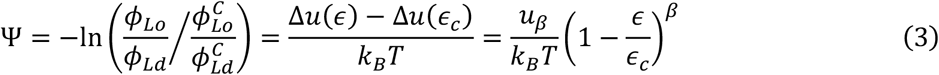

The concentration ratio was experimentally accessed through the measured fluorescence intensity ratio 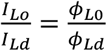 (Fig. 5A).

**Fig. 5:**
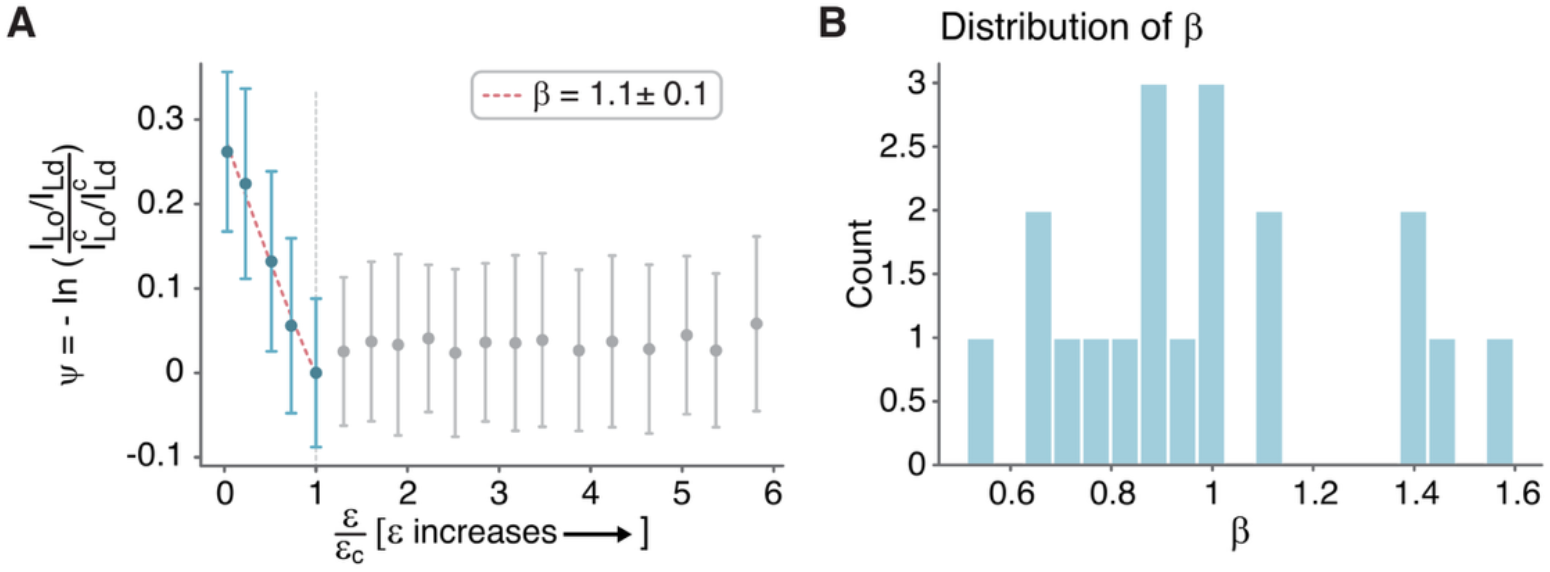
Fitting experimental strain-derived phase transition behavior to a power law theoretical model. **(A)** Order parameter Ψ vs. normalized strain *ε*/*ε*_*c*_ for a representative membrane. The order parameter vanishes at the critical point (*ε*/*ε*_*c*_ = 1). Red curve: power-law fit (Eq. 3) using the reduced strain, 1 − *ε*/*ε*_*c*_, yielding *β* = 1.1 ± 0.1 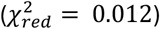. **(B)** Distribution of *β* values across all membranes (n=19) yielding *β* = 1.0 ± 0.3. Fit quality, 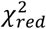, ranges from 0.001 to 1.099.

The order parameter Ψ decreased with increasing strain, exhibiting the expected power-law scaling relationship near the critical strain, and vanishing at the critical strain. Fitting was performed using the reduced strain, 1 − *ε*/*ε*_*c*_. However, for clearer visualization, data is presented vs. *ε*/*ε*_*c*_ such that applied strain increases along the horizontal axis. The red curve represents fitting using a single power-law scaling with two fitting parameters: the coefficient 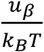 and the exponent *β*. Fitting across all membranes yielded *β* = 1.0 ± 0.3 (Fig. 5B) and 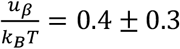. The strain-independent term, evaluated from the ratio at critical strain as 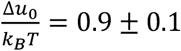.

The order parameter Ψ, constructed from the chemical equilibrium of Rh-PE between the Lo and Ld domains, exhibited robust power-law scaling with *β* ∼ 1 across all measured membranes and vanishing continuously at the critical strain. A single-term expansion successfully captured the observed behavior.

### A theoretical model successfully predicts *β*

To compare the experimentally observed exponent *β* with its theoretical expectation, we hypothesized that the strain-dependent free-energy difference Δ*u*(ε) (Eq. 1) originates from differences in the stretching energy of Rh-PE molecules between the Ld and Lo domains. In the absence of external stress, Rh-PE molecules adopt a given shape that differs from their preferred one. This difference depends on the local lipid environment of the probe, including molecular packing, local composition and substrate interactions, which differ between the domains. This mismatch is manifested as residual strain, denoted by 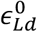 and 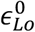 for the Ld and Lo phases respectively.

Upon applying external substrate strain *ϵ*, both domains experienced additional stretching, though not necessarily equally, as the strength of mechanical coupling to the substrate may differ between phases. We therefore describe the additional strain experienced by Rh-PE molecules in each domain as *α*_*Ld*\*Lo*_ · ∈, where the coefficients *α*_*Ld*\*Lo*_ represent the ratio between the externally applied strain and the actual local strain in each domain. The subscript *Ld* and *Lo* denotes the Ld and Lo phase. Based on these considerations, the elastic energy of a single Rh-PE molecule under external stretching compared to the stressless state is given by (25):

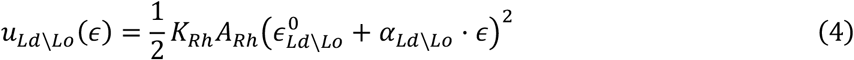

*K*_*Rh*_ and *A*_*Rh*_ are the area expansion-compression modulus of Rh-PE and its resting area per molecule. These are intrinsic properties of Rh-PE and are the same in both phases.

The energy difference related to strain, *u*_*Lo*_ − *u*_*Ld*_, is calculated using Eq. 4. To obtain the relation to the order parameter defined in the previous section, we insert this energy difference into Eq. 3, yielding the following general form for the order parameter:

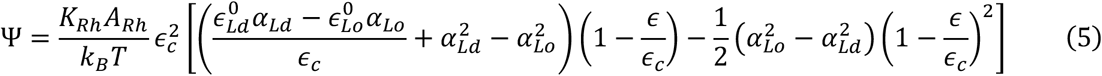

The coefficient of the linear term in Eq. 5 depends on the residual strains, 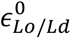, and differences in the membrane-substrate coupling, *α*_*Lo*/*Ld*_, whereas the quadratic term depends only on the coupling differences. If the coupling is the same in both domains (*α*_*Ld*_ = *α*_*Lo*_), as indicated by our area fraction analysis (Fig. S5C), Eq. 5 is reduced to

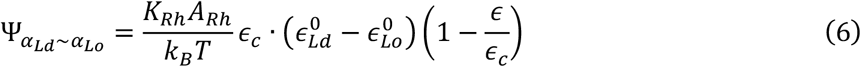

Hence, the scaling of the order parameter is linear with the reduced strain, *β* = 1, in agreement with the experimental observation of *β* = 1.0 ± 0.3. Thus, the linear scaling behavior near the critical strain is captured by the elastic model. The dominant contribution reflects the geometric mismatch between the intrinsic shape of Rh-PE and the local packing geometry imposed by the host domain, rather than differential strain between domains, which is comparable. At the critical strain, this geometric mismatch vanishes, and the Rh-PE distribution becomes uniform.

## Discussion

We investigated tension-mediated membrane reorganization in phase-separated SLBs subjected to controlled mechanical strain, using a motorized equibiaxial stretching device. Progressive substrate stretching induced membrane area strain, driving gradual homogenization of the Rh-PE distribution, with Lo and Ld domains becoming progressively less distinct, although slight residual partitioning persisted even above the critical strain.

Our results align with previous micropipette aspiration studies showing that membrane tension suppresses domain formation (15, 16). In those studies, tension was applied to membrane vesicles, leading to small decreases in the miscibility transition temperature; in contrast, we applied tension to supported bilayers under isothermal conditions. Despite differences in tension magnitude and experimental method, both approaches demonstrate tension-induced domain homogenization, consistent with theoretical predictions (17, 18). In contrast, studies in which membrane tension was induced via osmotic pressure reported the opposite behavior: osmotic shock promoted domain formation by increasing the miscibility transition temperature (12–14). These discrepancies may reflect distinct experimental conditions.

Several experimental limitations constrained our measurements. Photobleaching of the fluorescent probe limited the number of images acquired during each stretching experiment, resulting in limited sampling near the critical point where detailed information about scaling behavior is most critical. Image intensity normalization to correct for photobleaching was performed for presentation, and the analysis method accounts for photobleaching effects. However, this constraint limited the precision of critical exponent extraction. Since reduced *χ*^2^ is sensitive to the number of data points, the limited dataset may lead to overfitting, making quantitative comparison between alternative models challenging. Additionally, individual vesicles prepared from bulk lipid mixtures exhibit inherent compositional heterogeneity. This compositional variation gives rise to a broad distribution of miscibility transition temperatures (26). The observed distribution of critical strain values likely reflects this compositional variation rather than experimental noise.

Membrane hole defects appeared at higher strain levels, providing direct evidence of mechanical coupling between the bilayer and substrate. The exact strain at which initial holes formation occurred was difficult to determine due their small size; however, multiple visible holes were observed at the highest applied strains of 20-30%. These observations are consistent with the reported mechanical behavior of PDMS-supported lipid membranes, which begin to exhibit pore formation at strains of approximately 1–2% (27), yet remain stable during stretching experiments at 10–15% strain (28), significantly higher than the 1–5% lysis strain typical for unsupported biological membranes (29–31).

We have identified a tension-driven phase transition behavior from Lo–Ld coexistence to a homogeneous liquid phase through quantitative analysis of the fluorescence intensity ratio between Lo and Ld regions. This ratio increased progressively with area strain, exhibited distinct power-law scaling regimes before a well-defined critical strain, consistent with continuous phase transition behavior. The mean critical strain was 6 ± 3%. Using an area expansion-compression modulus of K_A_ ∼ 10^2^ mN/m (23, 32–34), this corresponds to a critical membrane tension on the order of ∼10^1^ mN/m. This estimate is in reasonable order-of-magnitude agreement with theoretical predictions by Uline et al (18).

We defined an order parameter Ψ based on Rh-PE probe equilibrium partitioning between domains, which vanished at the critical strain. Power-law fitting across all membranes yielded *β* = 1.0 ± 0.3, indicating near-linear scaling behavior. A simple elastic theory predicted *β* = 1 (Eq. 6). Beyond the critical strain, the membrane remained homogeneous upon further stretch, suggesting that the free energy transitions from two coexisting minima to a single minimum where phase coexistence disappears. As the membrane approached critical tension, the interfacial region between phases broadened, with domain boundaries becoming progressively less distinct. This broadening of the transition region, together with the shift between two minima of the free energy to one, is a characteristic of second order phase transition (24).

Physiologically, the dependence of the phase organization on membrane tension offers a potential mechanism for cellular regulation. The tension-induced suppression of domains could trigger or inhibit cell functions, allowing the cell to respond dynamically to mechanical stress.

## Materials and Methods

### PVA-assisted GUV formation

GUVs were prepared using a ternary mixture of saturated lipid (1,2-dioctadecanoyl-sn-glycero-3-phosphocholine (DSPC; PC(18:0)), unsaturated lipid (1,2-di-(9Z-octadecenoyl)-sn-glycero-3-phosphocholine (DOPC; PC(18:1)) and cholesterol in a 27/50/23 molar ratio. This composition was selected to induce coexistence of Lo and Ld phases at room temperature, with a DOPC-rich Ld phase and a DSPC-rich Lo phase, as previously characterized (22). For similar DSPC/DOPC/Chol compositions, miscibility transition temperatures of 35-38°C have been reported (20), ensuring phase coexistence at room temperature and near physiological conditions. 0.1% headgroup labeled fluorescent phospholipid 1,2-dioleoyl-sn-glycero-3-phosphoethanolamine-N-(lissamine rhodamine B sulfonyl) (Rh-PE) was added for membrane visualization. Bulky-headgroup Rh-PE fluorescent marker was used to visualize membrane microdomain formation and clearly distinguish between Lo and Ld phases, due to the preferential partitioning (22). All lipids were purchased from Avanti.

GUVs were prepared using a PVA-assisted swelling protocol (35, 36): 5 μl of lipid mixture at desired concentrations were applied to 25 mm PVA-coated coverslips and desiccated under vacuum for a minimum of 1 hour to ensure complete solvent removal. Vesicle formation was induced by adding 400 μl of growing buffer (70 mM NaCl, 25 mM Tris, 80 mM sucrose) and incubating at 50°C for 1 hour in a chamber. The resulting GUVs were collected and stored at 4°C with protection from light.

### SLB Formation on PDMS Substrates

PDMS substrates were activated using oxygen plasma (25% power, 0.4 mbar) for 50 seconds to make their surface more hydrophilic for vesicle rupture and bilayer formation (37). 70 μL of vesicle suspension was added within 1-2 minutes after plasma treatment, followed by immediate addition of 130 μL observing buffer (70 mM NaCl, 50 mM Tris, 55 mM Glucose) to prevent membrane exposure to air. After a 10-minute room temperature incubation, 10 μl of 2 mM CaCl_2_ was added to promote membrane adhesion to the substrate. Following an additional 10-minute incubation, the sample was carefully washed 10 times to remove unfused vesicles and debris and ensure a clean bilayer. Adhesion was evidenced by the occurrence of membrane rupture (holes) at substrate high stretching (SI Fig. S3).

### Fluorescence Microscopy

For observing phase separation in vesicles as shown in Fig. 1A, vesicles were placed in a 3D printed chamber filled with observation buffer and imaged using an Olympus IX73 inverted microscope with an X60 water immersion objective (NA = 1.1). For observing SLBs on a flexible substrate, as shown in Fig. 1C, imaging was done using an X40 objective (NA = 0.6) to enable a wider field of view with the same microscope system. SLBs on PDMS substrates were imaged directly within the custom-made PDMS chambers.

### Atomic Force Microscopy

#### Sample Preparation for AFM

To provide stability during SLB scanning, PDMS substrates were mounted on glass slides and activated using oxygen plasma as described above. Within 1-2 minutes after plasma treatment, 70 μL of vesicle suspension and 400 μL observation buffer were added. After 10 minutes, 10 μL of 2 mM CaCl_2_ in 600 μL buffer were added to promote membrane adhesion. Following another 10 minutes of incubation, samples were washed to remove unfused vesicles.

#### AFM Scanning

Nanoscale topography imaging was performed using the Bruker NanoWizard AFM system mounted on an Olympus IX73 inverted microscope, enabling correlative fluorescence-AFM imaging. MLCT-bio cantilevers (Bruker) with spring constants of 0.6 N/m (No. F) were used in intermittent contact mode (PeakForce™ mode) to minimize lateral forces on the samples and prevent tip damage. All AFM imaging was performed at room temperature (approximately 21°C) in buffer to maintain membrane hydration. Imaging parameters included setpoints of 300-600 pN, scan rates of 0.4-0.5 Hz, and resolutions of 128×128 or 256×256 pixels, depending on scan size. Direct Overlay™ enabled correlation between optical and AFM images. Fluorescence images of the scanned area were acquired, and AFM scanning was then performed on regions selected directly from the optical snapshots.

Images were processed using JPK Data Processing software. Processing included leveling and, when necessary, filtering or line replacement. Height profiles were extracted to quantify membrane thickness and inter-domain height differences.

### Equibiaxial stretching device and PDMS samples

A custom equibiaxial stretching device for controlled membrane tension experiments was constructed as previously described (38). In essence, it consists of six arms, made of stainless steel with CNC machining, that hold a PDMS sample connected via gears (SI Fig. S1A-B). Stretching movement was provided by a NEMA 17 stepper motor (Nanotec) with a planetary gearbox (Nanotec). The motor was controlled using a Tic 36v4 stepper motor controller (Pololu Robotics and Electronics) through the Tic Control Center software. The system enabled precise equibiaxial stretching compatible with a fluorescence microscopy system, achieving area strains of up to approximately 25% (see strain analysis below).

#### PDMS samples preparation

Polydimethylsiloxane (PDMS, Sylgard 184, Dow Chemical, US) samples consisting of a hexagonal frame bonded to a thin membrane substrate were fabricated for applying tension, based on the protocol established by Meyer et al.(38) The hexagonal frame bounds a central well (radius 1 cm) and was produced by mixing 4 g PDMS base with cross-linker (10:1 ratio), pouring into 3D-printed molds with an empty center cavity (SI Fig. S1C-D), degassing under vacuum for 10–20 min, and curing for 48 h at room temperature.

The supporting membrane (PDMS substrate) was fabricated in two layers on a silane-coated (1H,1H,2H,2H-Perfluorooctyltriethoxysilane, 97%, abcr GmbH, Germany) 3-inch silicon wafer. The base layer (4 g PDMS, 10:1 ratio) was spin-coated (150 rpm, 20 s; then 300 rpm, 40 s) to achieve ∼220 μm thickness, degassed, and cured for 35 min at 100°C. For calibration, 1.50 μm fluorescent beads (Fluoresbrite® Yellow Green Carboxylate Microspheres) were diluted 1:100 in isopropanol, vortexed, and sonicated for 30 min. The second layer (3 g PDMS, 10:1 ratio, with 200 μL beads solution) was spin-coated (750 rpm, 20 s; then 1500 rpm, 40 s) to create a ∼30 μm layer. The hexagonal frame was placed onto the uncured top layer with additional PDMS for adhesion, degassed, and cured for 3–4 h at 80°C. Access holes were punched and stabilized with PEEK sleeves.

#### Deformation evaluation and area strain calculation

Membrane deformation was evaluated using an image registration algorithm (ITK Elastix Python framework). This method quantified spatial transformations by performing pairwise mapping of reference and target images across the entire image. As the focal region shifts during stretching, images at all strain levels were manually aligned to the center to maintain a consistent field of view. Sequential microscopy images were registered pairwise using an affine transformation (AffineDTI).

The scaling factors *λ*_*x*_ and *λ*_*y*_ were extracted from the transformation parameters, representing the stretch ratio (*λ* = *L*/*L*_0_) in *x* and *y* directions. Strains were computed as *ε* = *λ* − 1.

The area stretch ratio can be calculated as (39)

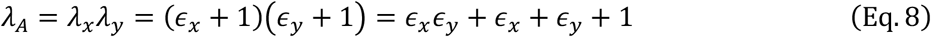

Area strain is

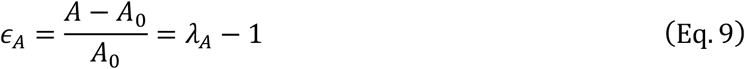

Therefore

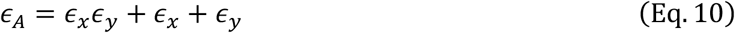

The stretching protocol achieved an area strain of approximately 25%, with equibiaxial deformation as presented in SI Fig. S2. Strain analysis was performed individually for each experimental sample to account for variability in device preparation and mounting.

### Membrane reorganization analysis using fluorescence intensity

Membrane homogenization was quantified by measuring local lipid composition through fluorescence intensity of Rh-PE, which serves as a proxy for the underlying lipid state (Lo/Ld). Low ratio values are expected when Rh-PE strongly favors the Ld phase in two-phase coexistence, while values approaching unity indicate more equal probe distribution, reflecting a transition toward uniform lipid composition across the membrane. Fluorescence intensity was quantified in each phase separately for each applied strain. Analysis was performed on non-normalized images; normalization was applied only for presentation purposes.

A region-based approach was used in which rectangular regions of interest were defined within the Lo and Ld phase domains, positioned away from domain boundaries to minimize artifacts. This approach was chosen over area-fraction analysis derived from whole-membrane intensity distributions, as large-area measurements were affected by optical artifacts introduced by debris in the buffer. The regional intensity ratios were computed from mean pixel values within those regions, providing values independent of the selected area dimensions and robustness against photobleaching artifacts (see SI Fig. S6C).

To estimate Lo and Ld phase area fractions, Multi-Otsu thresholding was applied to sequential fluorescence images to segment pixels into Lo and Ld phases at each stretching level. However, quantitative analysis of absolute phase areas was constrained by experimental limitations such as incomplete membrane imaging and residual buffer fluorescence, limiting accurate tracking of the total Lo/Ld area ratio across stretch levels (SI Fig. S4C). The segmentation was therefore used qualitatively, to assess changes in domain boundary sharpness and interfacial region width with increasing membrane tension.

To identify phase transition behavior from two phases to one phase, exponential functions were fitted to intensity ratio data. The critical strain was determined as the point where the exponential fit quality (*R*^2^) reached a maximum value, immediately before the intensity ratio plateaued. Membranes exhibiting two distinct exponential scaling regimes separated by a well-defined point (*ε*_*C*_) were classified as undergoing continuous phase transitions. Statistical analysis of critical strain values was performed across all vesicles exhibiting critical behavior.

### Model Fit Evaluation

Theoretical model fit quality was evaluated using the reduced chi-squared:

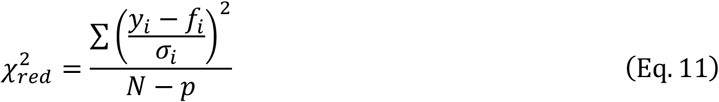

*N* is the number of data points, and *p* is the number of fitting parameters, *y*_*i*_ is the experimentally measured value, *f*_*i*_ is the model value and *σ*_*i*_ is the error of *y*_*i*_.

## Supporting Information

Additional experimental details and methods (DOCX).

## Acknowledgments

A.P.I., G.G., and R.S thank Shigeyuki Komura and David Andelman for helpful discussions on the theory. Code and text were refined with the support of AI tools.

R.S. acknowledges support by the Israel Science Foundation (Grants No. 1289/20 and 2303/25) and the NSF-BSF (grant no. 2021793). Co-Funded by the European Union (ERC ReMembrane 101077502). Views and opinions expressed are, however, those of the authors only and do not necessarily reflect those of the European Union or the European Research Council Executive Agency. Neither the European Union nor the granting authority can be held responsible for them. S.K. acknowledges funding from the German Research Foundation (Deutsche Forschungsgemeinschaft, DFG) in the framework of projects 430255655 (KO 3572/8-1) and 449544493. G.G. acknowledges support by the Israel Science Foundation (grant no. 913/26).

## Author contributions

A.P.I., G.G., and R.S. designed the research, and A.P.I. performed the research and data analysis. R.M. and S.K. designed the equibiaxial stretching device and compatible PDMS sample geometries and provided the design files and component specifications. A.P.I. built a local version of the device. G.G. and R.S. contributed to data analysis and interpretation. A.P.I. and G.G. developed the theoretical framework. R.S. supervised the research, and A.P.I., G.G., and R.S. wrote the paper.

## Notes

### Competing Interest Statement

The authors have declared no competing interest.

### Summary of Updates

Graphics in figures 1-5 have been updated, and new illustrations have been added for clarification.

