## Supplementary Information for "Stretching drives Membrane Homogenization in Phase-Separated Supported Lipid Bilayers"

### 1. Equibiaxial stretching assay

#### 1.1. Equibiaxial stretching device and PDMS samples

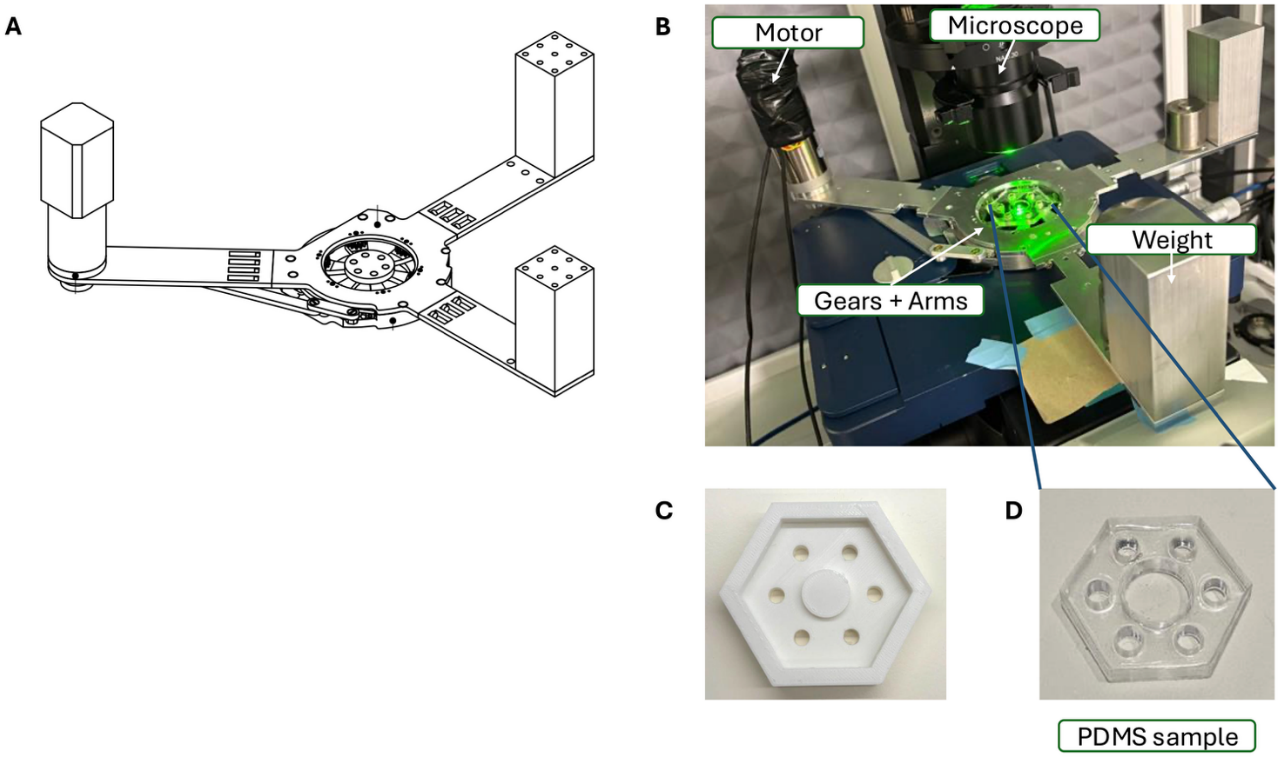

**Fig. S1:** To apply controlled strain to the SLBs, a custom-designed motorized equibiaxial stretching device was constructed. **(A)** Design of the custom motorized biaxial stretching device. **(B)** Image of the implementation of the stretching device mounted on a microscope stage. **(C)** 3D-printed mold used to create hexagonal PDMS substrates for uniform tension application. **(D)** The PDMS layers were fabricated into samples with a designated shape compatible with the equibiaxial stretching device, formed using 3D-printed molds. Hexagonal PDMS substrate with a central well for SLB formation.

#### 1.2. Membrane area strain under equibiaxial tension

##### *1.2.1. Membrane area strain evaluation*

**A**

**B**

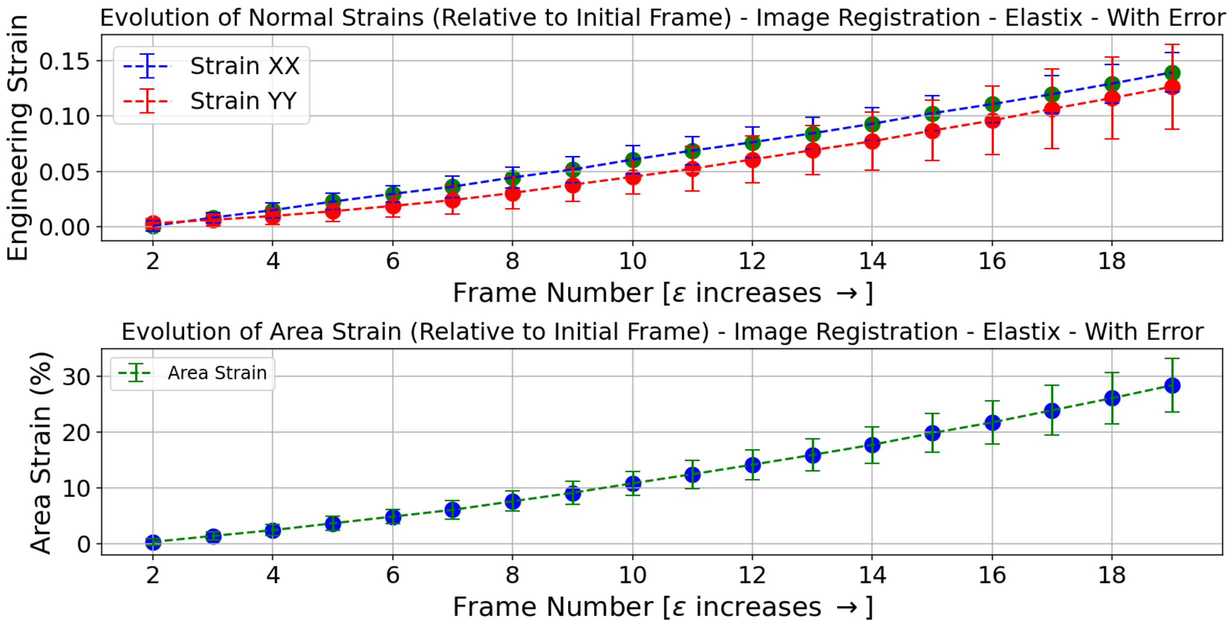

**Fig. S2: Deformation Analysis of membrane strain during equibiaxial stretching. (A)** Evolution of normal strains along the X-axis (blue) and Y-axis (red) as a function of frame number during progressive stretching. The system exhibited minor anisotropy between axes while maintaining consistent equibiaxial deformation throughout the experiment. **(B)** Evolution of total area strain during the stretching protocol, reaching approximately 28% at the final frame. Strain values were quantified using image registration analysis (Elastix). The progressive increase in area strain confirmed that supported lipid bilayers were mechanically coupled to the substrate and subjected to tension during deformation.

##### *1.2.2. Error Estimation of membrane area strain evaluation*

To quantify the uncertainty associated with area strain measurements derived from image registration, an error estimation procedure based on multi-partition sub-region analysis was used. For each frame pair, the image was divided according to three spatial partition schemes (2×2, 3×1, and 1×3 grids). Within each partition, affine image registration was performed independently on each sub-region using Elastix. The resulting scale parameters were used to compute incremental stretch values in the x and y directions, from which cumulative engineering area strain was derived for each sub-region as described in Methods.

The representative area strain for each partition was taken as the median of the cumulative area strains of the sub-regions, excluding sub-regions identified as outliers due to objects drifting across sub-region boundaries. The final error estimate for each frame was computed as the sample standard deviation across the three partition medians.

##### 1.3. Membrane stretching on flexible substrate using equibiaxial stretching device: rupture as evidence of mechanical coupling

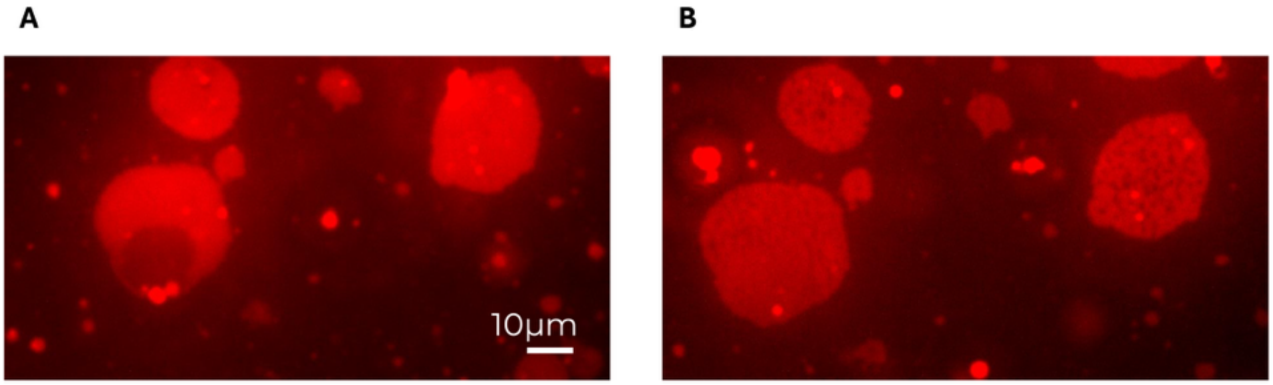

**Fig. S3: Membrane defects induced by high tension. (A)** Membrane at low strain showing intact bilayer structure. **(B)** Membrane at maximal area strain (~23.81%) displaying tension-induced visible holes (dark circular defects with lower fluorescence intensity than the surrounding membrane). The appearance of membrane defects at high strain levels provided direct evidence of mechanical coupling between the bilayer and substrate, confirming that SLBs were subjected to tension rather than flowing across the substrate surface during deformation.

2. Representative experiment: Complete image sequence across all applied tension levels showing progressive membrane homogenization

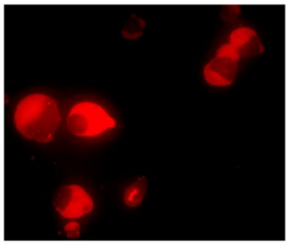

$\Delta A = 0$

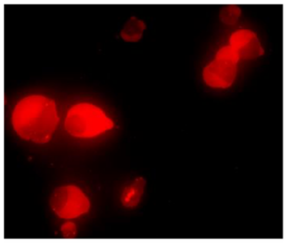

$\Delta A = 0.15\%$

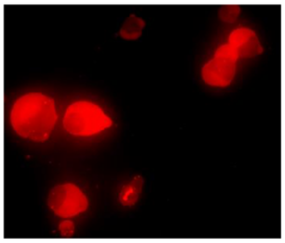

$\Delta A = 1.11\%$

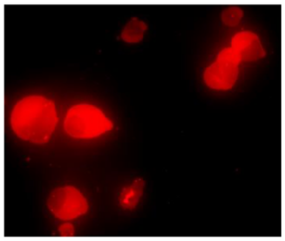

$\Delta A = 2.48\%$

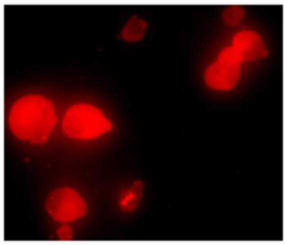

$\Delta A = 3.51\%$

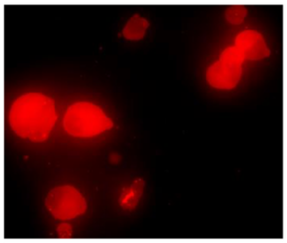

$\Delta A = 4.82\%$

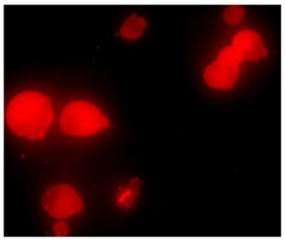

$\Delta A = 6.27\%$

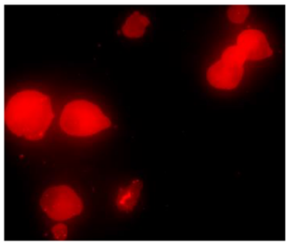

$\Delta A = 7.74\%$

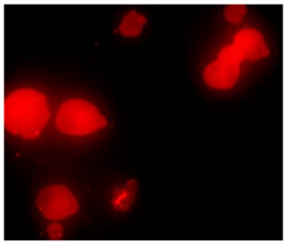

$\Delta A = 9.14\%$

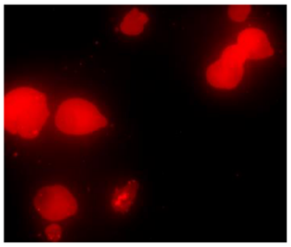

$\Delta A = 10.74\%$

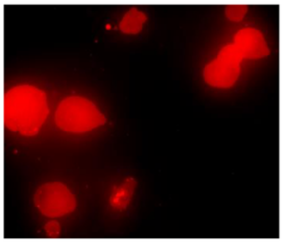

$\Delta A = 12.15\%$

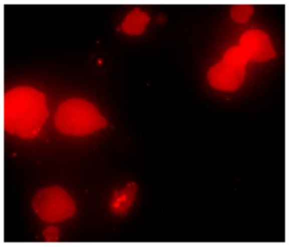

$\Delta A = 13.74\%$

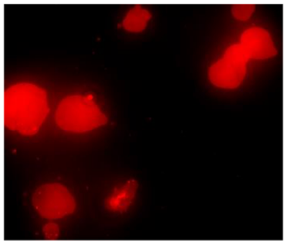

$\Delta A = 15.32\%$

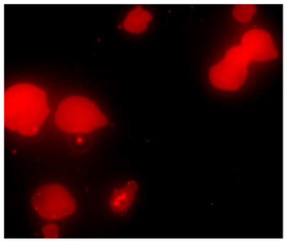

$\Delta A = 16.74\%$

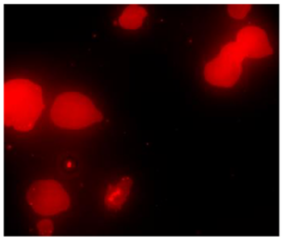

$\Delta A = 18.66\%$

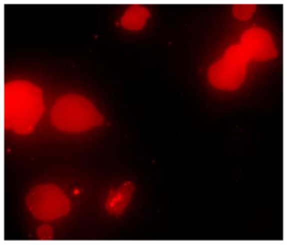

$\Delta A = 20.42\%$

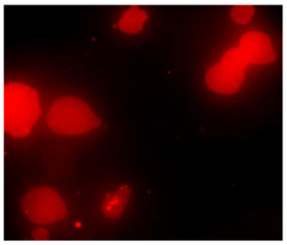

$\Delta A = 22.37\%$

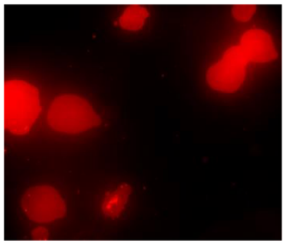

$\Delta A = 24.34\%$

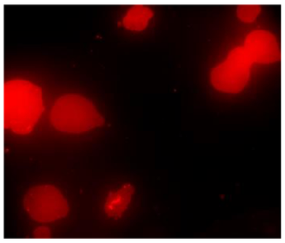

$\Delta A = 25.88\%$

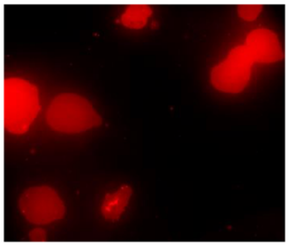

$\Delta A = 28.01\%$

**Fig. S4: Representative fluorescence microscopy images of phase-separated lipid membranes labeled with Rh-PE at sequential imposed area strains.** Images from $\epsilon = 0$ (unstrained) to $\epsilon = 0.28$, demonstrating the progressive evolution of Lo (dark) and Ld (bright) phase domains. At low strains, distinct phase domains with sharp boundaries were observed. Intermediate strains exhibited gradual domain boundary diffusion and reduced fluorescence contrast between phases, indicative of partial lipid mixing. At high strains ($\epsilon > 0.05$), domains became indistinguishable, with near-uniform fluorescence distribution consistent with mechanical homogenization of the membrane. Image intensities were normalized to account for photobleaching effects during presentation.

#### 3. Quantitative analysis of membrane organization under tension

##### 3.1. Domain area-fraction evaluation

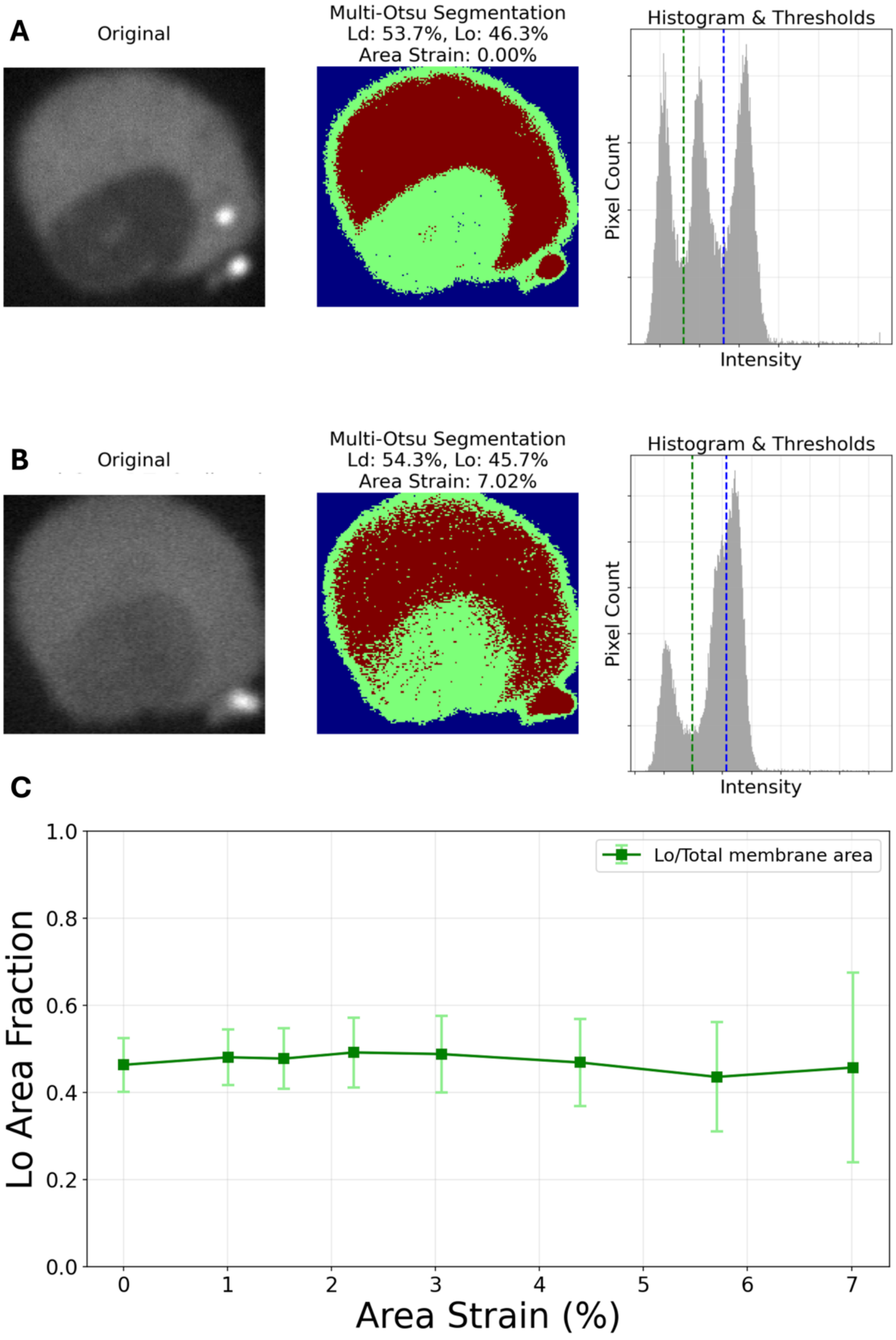

**Figure S5. Phase area fraction analysis under membrane tension. (A-B)** Representative fluorescence images (left) and their corresponding Multi-Otsu segmentation (middle) at area strains of 0% **(A)** and 7.02% **(B)**, immediately before the critical strain. Pixels were classified into Lo (dark, green) and Ld (bright, red) phases, with background pixels excluded (blue). Intensity histograms (right) show pixel intensity distributions with the Multi-Otsu thresholds (dashed lines) used to define phase boundaries at each stretching level. Incomplete membrane imaging and residual buffer fluorescence limited absolute phase area quantification. However, domain boundaries visibly broadened and became less distinct with increasing membrane tension. **(C)** Lo area fraction (Lo/total membrane area) as a function of area strain for the same membrane shown in (A-B). Error bars represent segmentation uncertainty estimated from threshold variations. The large and increasing error bars at higher strain levels reflect the difficulty of reliably area evaluation of Lo and Ld phases. Under these measurement constraints, the ratio of Lo to Ld phase areas appeared approximately constant before the critical strain, illustrating the limitation of area fraction quantification as a measure of membrane homogenization under tension.

#### 3.2. A region-based approach to evluate Rh-PE fluorescence intensity ratio between domains

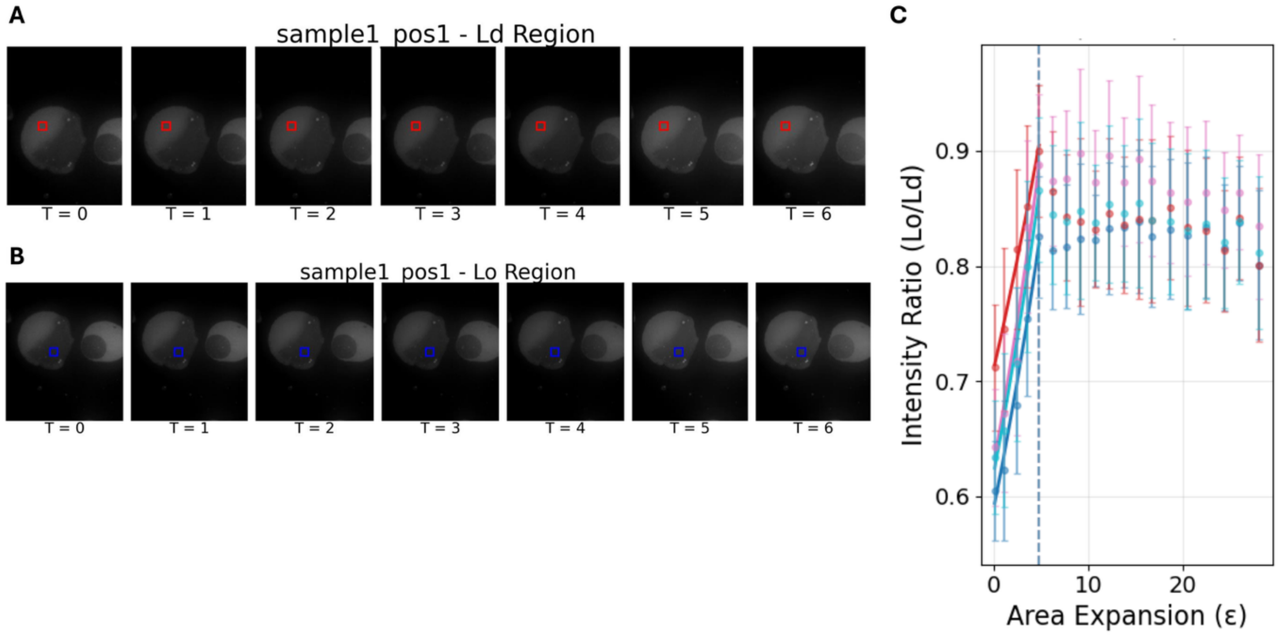

**Figure S6. (A-B)** A region-based approach was used in which rectangular regions of interest were defined within the Ld **(A)** and Ld **(B)** phase domains, positioned away from domain boundaries to minimize artifacts. **(C)** The regional intensity ratios were computed from mean pixel values within those regions, providing values independent of the selected area dimensions and robustness against photobleaching artifacts. Error bars represent the standard deviation of the mean values across the selected regions. All the graphs have taken from the same membrane while choosing different regions of interest, and they all within each other errors, yielding the same critical strain.

#### 4. Order parameter framework

In order to analyze a phase transition near to a critical point, we defined an order parameter $\Psi$. The order parameter was constructed based on chemical potential equilibrium requirements. In the two-phase coexistence state, the Rh-PE probe energy in Ld and Lo domains maintains a chemical potential equilibrium:

$$\mu_{Ld} = \mu_{Lo} \tag{S1}$$

The chemical potential in each region is defined by the local concentration of Rh-PE (number of Rh-PE molecules per unit area in the Ld/Lo phase):

$$\begin{aligned} \mu_{Ld} &= \mu_{Ld}^0 + k_B T \cdot \ln\left(\frac{\phi_{Ld}}{1-\phi_{Ld}}\right) \\ \mu_{Lo} &= \mu_{Lo}^0 + k_B T \cdot \ln\left(\frac{\phi_{Lo}}{1-\phi_{Lo}}\right) \end{aligned} \tag{S2}$$

Where $\phi_{Ld}$ and $\phi_{Lo}$ represent the concentration of Rh-PE in the Ld and Lo domains, respectively. Note that $\phi_{Ld} + \phi_{Lo} \neq 1$. Substituting in chemical potential equilibrium (Eq. S1):

$$\ln\left(\frac{\phi_{Ld}}{1-\phi_{Ld}}\right) + \frac{\mu_{Ld}^0 - \mu_{Lo}^0}{k_B T} = \ln\left(\frac{\phi_{Lo}}{1-\phi_{Lo}}\right) \tag{S3}$$

Since Rh-PE is in small total concentration (0.1%) we expand Eq. S3 in $\phi_{Lo}, \phi_{Ld} \ll 1$

$$\ln\left(\frac{\phi_{Ld}}{\phi_{Lo}}\right) = -\frac{\Delta u}{k_B T} \tag{S4}$$

With $\Delta u = \mu_{Ld}^0 - \mu_{Lo}^0$. Since the energy difference close to the critical point is small, we expanded $\Delta u$ in a power series around the critical point

$$\Delta u \sim u_0 + u_1 \left(1 - \frac{\epsilon}{\epsilon_C}\right)^{\beta} \tag{S5}$$

The order parameter was normalized by the critical value (where $\epsilon = \epsilon_C$). Experimentally, this concentration ratio was accessed through the measured intensity ratio $\frac{I_{Lo}}{I_{Ld}}$, which yielded the following order parameter:

$$\Psi = -\ln\left(\frac{I_{Lo}}{I_{Ld}} \Big/ \frac{I_{Lo}^C}{I_{Ld}^C}\right) \tag{S6}$$

This normalized quantity vanishes at the critical point.
